# Deep convolutional and recurrent neural networks for cell motility discrimination and prediction

**DOI:** 10.1101/159202

**Authors:** Jacob C. Kimmel, Andrew S. Brack, Wallace F. Marshall

## Abstract

Cells in culture display diverse motility behaviors that may reflect differences in cell state and function, providing motivation to discriminate between different motility behaviors. Current methods to do so rely upon manual feature engineering. However, the types of features necessary to distinguish between motility behaviors can vary greatly depending on the biological context, and it is not always clear which features may be most predictive in each setting for distinguishing particular cell types or disease states. Convolutional neural networks (CNNs) are machine learning models allowing for relevant features to be learned directly from spatial data. Similarly, recurrent neural networks (RNNs) are a class of models capable of learning long term temporal dependencies. Given that cell motility is inherently spacio-temporal data, we present an approach utilizing both convolutional and long- short-term memory (LSTM) recurrent neural network units to analyze cell motility data. These RNN models provide accurate classification of simulated motility and experimentally measured motility from multiple cell types, comparable to results achieved with hand-engineered features. The variety of cell motility differences we can detect suggests that the algorithm is generally applicable to additional cell types not analyzed here. RNN autoencoders based on the same architecture are capable of learning motility features in an unsupervised manner and capturing variation between myogenic cells in the latent space. Adapting these RNN models to motility prediction, RNNs are capable of predicting muscle stem cell motility from past tracking data with performance superior to standard motion prediction models. This advance in cell motility prediction may be of practical utility in cell tracking applications.

## I. Introduction

CELL motility is an emergent property of living matter that spans the nanomolecular and macroscopic length scales, involving a complex regulatory network and dynamic reorganization of the cell’s geometry [1], [2]. Cells can display a diverse set of motility behaviors, and these behaviors can provide a useful window for inference of a cell’s functional state. Neoplastic transformation has long been appreciated to alter cell motility behaviors, increasing the migration rate of various models in culture and serving as a mechanism for metastasis [3], [4], [5], [6], [7]. The motility behaviors of cancer cells in culture can even be predictive of broader tumor progression [8].

Likewise, the migration of progenitor cells is critical in early development and tissue regeneration [9]. Skeletal muscle stem cells (MuSCs) provide an accessible cell culture system to study stem cell motility behaviors *in vitro* by timelapse imaging. During embryonic development, MuSC precursors must migrate from early stage developmental structures (somites) to their adult location along the edge of muscle fibers in the trunk and limbs [10], [11]. In the adult, motility continues to play a critical role, as MuSCs migrate along muscle fibers *in vivo* to sites of injury to initiate tissue repair [12], [13]. Motility behaviors are heterogeneous between MuSCs and change during stem cell activation [14], [15]. Heterogeneous fitness for regeneration within the MuSC pool is well appreciated [16], and analysis of heterogeneous motility behaviors may provide an additional lens through which to decompose different MuSC phenotypes.

Given the biological importance of motility behaviors, classification of cells based on motility behaviors has useful applications in research and diagnostics. Similarly, exploration of heterogeneity within the motility behaviors of a cell population may provide biological insights. However, it is often difficult to determine which features of motility behavior will be predictive of a phenotype of interest, or allow for discrimination of heterogeneous behavior. Different phenotype classification tasks and cell populations may require distinct feature sets to extract valuable biological information. A method to algorithmically determine relevant features of cell motility for a given classification or discrimination task is therefore advantageous.

### A. Related Work

To date, a number of tools have been proposed that rely upon a set of handcrafted features to quantify cell motility behaviors, providing some remarkable results [17], [18], [19], [20], [21]. Neural progenitor cells were discriminated by morphology and motility behavior alone [21], and genes that affect motility have been identified solely from timelapse imaging data [17]. Similarly, a heuristic cell motility feature was identified as one of the most important sources of information to predict hematopoetic cell lineage decisions [22]. We have recently demonstrated that rates of cell state transitions and the ordered or random nature of these transitions may also be inferred from motility alone [15].

These results demonstrate the potential insights that may be gathered from more extensive analysis of cell motility. However, these methods rely upon engineering of a handcrafted feature set, and have thus far focused largely on features of speed and directional persistence. It is possible that more complex features may allow for improved discrimination of cell motility behaviors, but it is difficult to predict what these features may be in each context.

Convolutional neural networks provide an approach to learn relevant features from data, rather than handcrafting features based on a “best guess” of which features are relevant. In the field of computer vision, convolutional neural networks (CNNs) have recently made rapid advancements, demonstrating state-of-the-art performance on a variety of image classification tasks [23], [24], [25], [26]. CNNs utilize a set of parametrized kernels to extract spatial features, allowing distinct feature kernels to be learned for a given classification task [27]. In this way, CNNs are able to learn a “representation” of the problem’s feature space. Feature space representations may also be learned in an unsupervised manner by training CNN autoencoder architectures to encode and decode [28], [29]. This approach may be useful for learning relevant motility features where an explicit classification task is not present.

While CNNs are most commonly applied to tasks involving analysis in images with two spatial dimensions and one channel dimension at a single time-point, convolution is a natural analytical tool for any input information with spatial dimensions. CNNs have been successfully applied to a diverse set of non-imaging domains, including natural language processing [30], bird song segmentation [31], and EEG recordings [32]. Perhaps most clearly mirroring our challenge of motion classification, CNNs have performed well in the classification of video recordings [33], [34], [35], [36].

Cell motility may be represented as a time series with one spatial dimension and one channel dimension, where the channel dimension represents the Cartesian axes of motion (i.e. *x*, *y*) and the spatial axis represents time. In this formulation, the value of the *t*, *c* element in the (Time, Coordinate) matrix represents the position value for a tracked cell at time *t* in coordinate *c*. Multiple problem domains have shown success in applying CNNs to multi-channel time series data in this manner [37], [38], [39], [40]. Similarly, convolutional layers with one spatial dimension and one channel dimension may allow for motility behavior classification and unsupervised feature learning without *a priori* definition of handcrafted features.

Deep neural networks have also been extensively applied to the analysis of sequential inputs, such as natural language sentences and biological polymer sequences [30], [41], [42], [43]. While simple 1D CNNs that consider raw sequence inputs can be effective, the introduction of recurrent units such as long short-term memory (LSTM) units to learn temporal relationships within the input sequence can improve performance and effectively learn long-term dependencies [44]. In the multi-channel time series representation described above, CNN layers may function as feature extractors that require no hand-engineering.

Pairing these extracted features with recurrent neural network (RNN) units may also allow for motility behavior analysis capable of learning long-term dependencies across the time series. Similar approaches combining convolutional layers and recurrent units have proven effective in the analysis of biological polymer sequences [45] and in image classification [46].

Here, we investigate whether RNNs paired with CNN feature extractors could be effectively applied to the problem of cell motility behavior classification. We develop a tool we call *Lanternfish* to represent motility paths as multi-channel time series, classify different motility behaviors, learn motility features in an unsupervised fashion, and predict future cell motility from past behavior using RNNs. *Lanternfish* represents cell motility as a simple multi-channel time series, where time series values are Cartesian coordinates. We demonstrate that RNNs with convolutional layers as feature extractors are sufficient to accurately distinguish experimentally observed cell motility behaviors. Autoencoder architectures based on these models capture variation between cell types and reveal heterogeneity within cell states in the latent space. Additionally, we show that our RNN model can be adapted to predict cell motility in subsequent frames more accurately than standard methods, with potential applications in the field of cell tracking.

## II. Methods

All implementations for work presented here are available on Github at https://github.com/jacobkimmel/lanternfish. Experimentally measured cell motility data and cell motility simulators are available in https://github.com/cellgeometry/heteromotility.

### A. RNN Classification Architectures

Our baseline RNN classification architecture utilizes an LSTM layer with *n* = 256 hidden units, followed by two fully-connected layers with 256 and 128 units respectively. Each of these fully-connected layers is paired with a rectified linear unit activation [23]. The final layer is a fully-connected layer with *n* = *K* units where *K* is the number of classes paired with a softmax activation. This baseline architecture is presented in Fig. 1A.

**Fig. 1.**
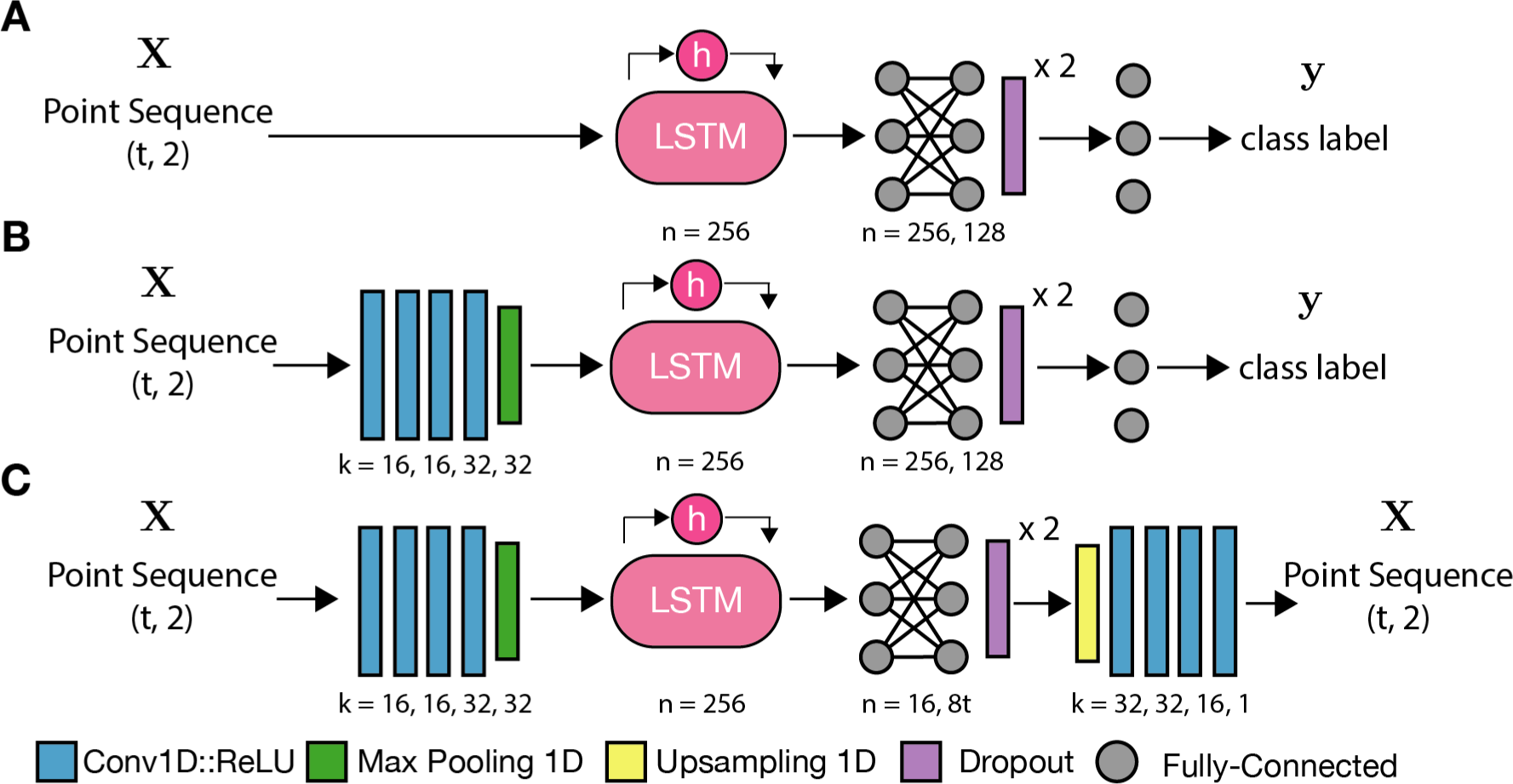
Cell motility classification and autoencoder architecture schematics. (A) A baseline RNN architecture, without convolutional feature extractors. (B) RNN classification and (C) autoencoder architectures with convolutional feature extractors, where *k* = *i, j*, … is the number of parameterized kernels used by each 1D convolutional layer in a series, and *n* = *i, j*, … is the number of nodes in a fully-connected layer or LSTM unit in a series. Convolutional layers and paired with a rectified linear unit activation. Pooling and upsampling layers operate with isotropic kernels of size 2 and stride of 2. Zero padding is performed as needed in autoencoder models to match input size.

We modify this architecture slightly to yield an RNN with convolutional feature extractors. This architecture prepends a set of four convolutional layers with one spatial dimension and one channel dimension (“1D” convolutions for brevity). These 1D convolutional layers utilize size 3 kernels with unit strides. Following these convolutional layers is a max pooling layer with filter size 2 and stride *s* = 2. Following the max pooling layer, the second architecture is identical to the first (Fig. 1B).

### B. RNN Autoencoder Architecture

Our RNN autoencoder architecture strongly resembles the classification network. Following the fully-connected layers in the classification architecture, the RNN autoencoder appends a 1D upsampling layer and mirror 1D convolutional layers to return the input back to the original size (Fig. 1C). Mean-squared error (MSE) against the input sequence was utilized as a loss function for training.

### C. RNN Motility Prediction Architecture

We adapt our RNN autoencoder architecture to a prediction architecture by removing the max pooling, fully-connected, and dropout layers. Sequences are convolved by four 1D convolutional layers, as in the autoencoder, before being passed to an LSTM and convolved by four more 1D convolutional layers. The final convolutional layer uses a linear activation function rather than a ReLU. Input sequences length *τ_in_* are provided in the same multi-channel time series format as our other RNN architectures, and output sequences are multi-channel time series of length *τ_out_*. The number of LSTM units is adjusted to *n* = 2*τ_out_* depending on the length of desired output sequences.

### D. Baseline Motility Classification

As a baseline motility classifier, a heuristic feature extractor is paired with a Random Forest (RF) classifier [47]. The feature extractor calculates 13 parameters of motion: (1) mean displacement, (2) displacement variance, (3) minimum and (4) maximum displacement, (5) the mean turning angle, (6) the mean turning angle magnitude, (7) turning angle variance, (8) total distance traveled, (9) net distance traveled (distance from starting position to final position), (10) progressivity of motion (net distance as a fraction of total distance) [48], (11) linearity of motion (Pearson’s *r*^2^), (12) monotonicity of motion (Spearman’s *ρ*), and (13) the convex hull area of the cell motility track [17]. These heurstics are commonly employed in the quantitative cell motility literature [21], [48], [49], [17]. The RF classifier hyperparameters were optimized by grid search for each application. Code for the baseline classifier is available on Github.

### E. Baseline Kinematic Motion Prediction

A linear kinematic model is used for baseline motility predictions. The kinematic model calculates the mean velocity

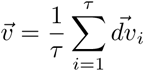

across the last *τ* time steps in the preceding track and projects the object by 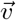 for each predicted time step. The temporal window *τ* is optimized by parameter search. This model assumes the moving particle exhibits ballistic motion.

### F. Saliency Analysis

The “saliency” of regions within an input track, estimating their importance for the assignment of a specific class by a classification model, was performed in the manner described previously [50]. Briefly, for a given input track *x*, class *c*, and trained trained classification model *f* (*x*) that outputs the class score for *c*, *f_c_*(*x*), each element in *x* is evaluated for influence on the score *f_c_*(*x*). This influence is computed as the gradient on *x* with respect to the class score *f_c_*(*x*). We present the rectified gradients as saliency maps using a ReLU.

### G. Computational Infrastructure

Nvidia GTX 1080 (Pascal) and Titan Xp GPUs were used for all experiments.

### H. Cell Culture

Mouse embryonic fibroblasts, muscle stem cells, and myoblasts were cultured as previously described [15]. Neoplastic MEFs were generated as described and generously donated by the authors of [51].

### I. Timelapse Cell Imaging

Timelapse cell imaging, cell segmentation, and cell tracking was performed as described [15]. Briefly, cells were imaged for 10 hours in DIC at 6.5 minute intervals using a stagetop incubator at 37°C and 5% CO_2_. Images were segmented using common heuristic techniques and tracking was performed using a modified version of uTrack [52]. Cell tracking data is available on the “Heteromotility” Github repository https://github.com/cellgeometry/heteromotility.

## III. Experimental Results

### A. Motility Simulations

To determine if RNN models could discriminate between different types of motion under ideal conditions, we trained RNN classification models on simulated data from 3 distinct, biologically relevant models of motion, namely random walks, Levy flights, and fractional Brownian motion. Random walks are a type of motion with normally distributed random step sizes and directionality. Random walks are observed in freely diffusing biomolecules [53]. Levy flights similarly display random directionality, but step sizes are instead chosen from a long-tailed Levy distribution. Levy flights are observed in multiple biological systems and optimize path finding [54], [55], [56], [57]. Fractional Brownian motion models a random walk with long term dependence, similarly relevant as a representation of regulated motion in biology [58], [59]. By starting with simulated data we can optimize parameters using large sample sizes that would be difficult to obtain with living cells.

Random walks, Levy flights, and fractional Brownian motion were simulated for classification, each with a mean displacement of *µ* = 5 (*x, y*) units per time step. Simulations were carried out for *T* = 100 time steps and restricted to a (2048, 2048) pixel plane, representing the field-of-view that might be expected using a standard 4 megapixel microscopy camera. Example simulations are presented alongside experimentally measured cell motility tracks (Fig. 2).

**Fig. 2.**
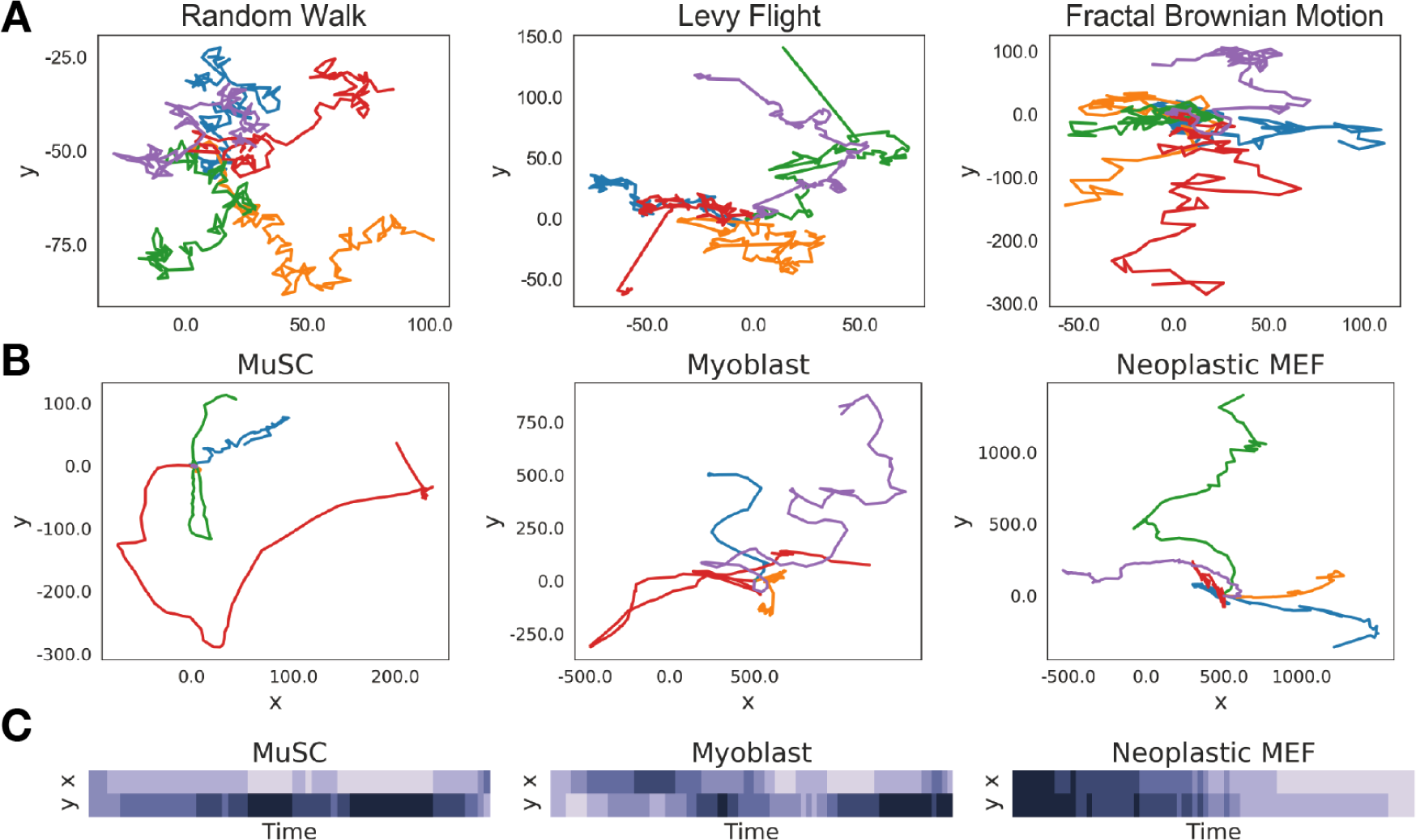
(A) Representative motility simulations of each class. (B) Representative cell motility tracks for different cell types. (C) Heatmaps demonstrating the mutli-channel time series representation of cell motility for representative tracks of multiple cell types.

### B. RNNs accurately classify simulated motility behaviors

Experiments to classify multi-channel time series representations of cell motility were performed using *n* = 15, 000 samples from each simulation class. RNN classifiers were trained and evaluated using 5-fold cross validation. In each training fold, 10% of the data was used for testing and model selection by early stopping. Early stopping was performed in all models after the testing loss failed to improve for 3 consecutive epochs [60]. Models were evaluated based on the prediction accuracy on the validation set.

Models were trained using crossentropy as a loss function. Optimization was performed with *Adam* using a learning rate of *ϵ* = 0.001 [61]. Weights for each of the convolutional and fully-connected layers were regularized by the *l*_2_ norm with strength *λ* = 10^−6^. The optimizer, learning rate, and regularization strength were chosen empirically, and a rigorous search for optimization hyperparameters was not performed. Using this scheme, models fit rapidly on simulated data (Fig. 3A).

**Fig. 3.**
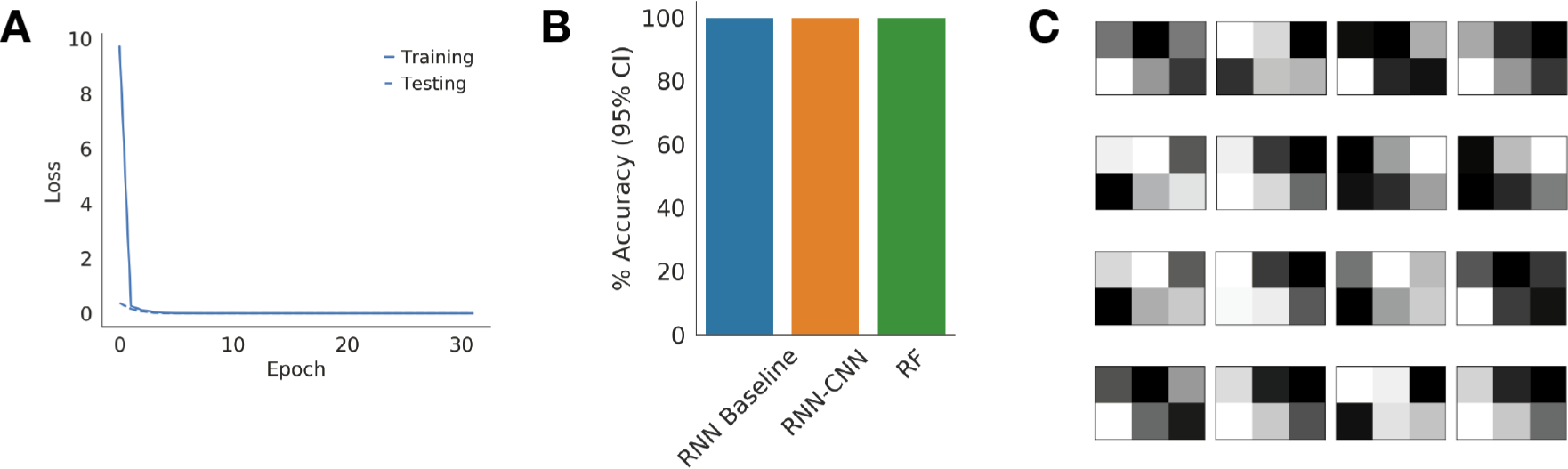
RNN classifiers effectively distinguish simulated models of motion. (A) Representative training curves for a baseline RNN classifier and an RNN classifier with convolutional feature extractors. (B) Performance of a baseline RNN classifier, RNN classifier with convolutional feature extractors, and a Random Forest (RF) baseline trained on heuristic features. Bars presented are the mean accuracy ±95% confidence intervals. (C) Kernel weights learned in the first convolutional layer of the RNN model diagrammed in Fig. 1B.

Both baseline RNN models and RNN models with convolutional feature extractors achieved near perfect classification accuracy (Fig. 3B). Likewise, a Random Forest model trained on hand-engineered features successfully disciminated these simulations. This result indicates that RNNs are capable of discriminating different types of motion with no hand-engineering of features.

To investigate how the RNN models may be learning to discriminate different motion models, we visualized filters from the first convolution layer of RNNs with convolutional feature extractors (Fig. 3C). Inspection of these filters reveals that several combinations of gradients are learned across the first and second dimension of motility. Interpreting filters in later layers becomes more challenging, but this result is in line with intuitions and suggests that at least some features we may engineer by hand (metrics of displacement, etc.) are being learned by convolutional feature extractors.

### C. RNNs accurately discriminate cell types by motility behavior

After validating that RNNs were sufficient to distinguish simulated classes of motion, we applied the same classification networks to distinguish different types of experimentally measured cell motility. Cell motility was tracked in three different cell types by timelapse imaging for 10 hours, followed by segmentation and tracking by standard methods. Mouse embryonic fibroblasts (MEFs) are commonly used for *in vitro* cell culture assays, and neoplastic transformation of these cells has been demonstrated to alter their motility behaviors [15]. We tracked both wild-type and neoplastic (*c-Myc* overexpression, *HRas-V12*) MEFs to compare their motility behaviors. Muscle stem cells (MuSCs) are the obligate stem cell of the skeletal muscle, and their motility is known to be effected by their activation state [14]. Activated MuSCs commit to become myoblasts, a proliferating myogenic progenitor cell. We tracked both MuSCs and myoblasts to compare motility between these states of myogenic commitment (see Methods for culture details).

To determine if RNNs could distinguish cell types based on experimentally measured motility, we trained RNN classifiers to discriminate between MEFs and MuSCs. Models were trained for 1000 epochs using early stopping with a 15 epoch patience period. Training was performed on a total of *n* = 562 MuSC and *n* = 562 MEF tracks (both wild-type and neoplastic) using 5-fold cross validation. Equal numbers of MuSCs and MEFs were used to ensure class balancing. In each training fold, 10% of the data was reserved for testing during the training process and model selection.

In other domains, transfer learning between models trained on different data sets has proven advantageous to model performance [62]. It is possible that pre-training an RNN classification model on the simulated motility tracks described above, where training data is plentiful, will yield performance improvements when classifying experimentally measured cell motility, where data is expensive. To assess this possibility, we performed the classification experiments above both with random initializations (*de novo* training) and with weights transferred from a corresponding model trained to classify simulations (Simulation pretraining).

Mean validation accuracy on this cell type classification task was ∼ 85% for baseline RNN models and ∼ 95% for RNN models with convolutional feature extractors when trained *de novo*, compared to a heuristic RF baseline model at ∼ 95% (Fig. 4A). These results indicate that even with a small data set such as this, RNN models can be effectively trained to discriminate different types of cell motility with performance similar to hand-engineered features (Fig. 4A). Moreover, RNN models with convolutional feature extractors improve significantly upon the RNN baseline (*p* < 0.01, *t*-test), suggesting that convolutional layers allow the models to capture features that are missed by LSTMs alone.

**Fig. 4.**
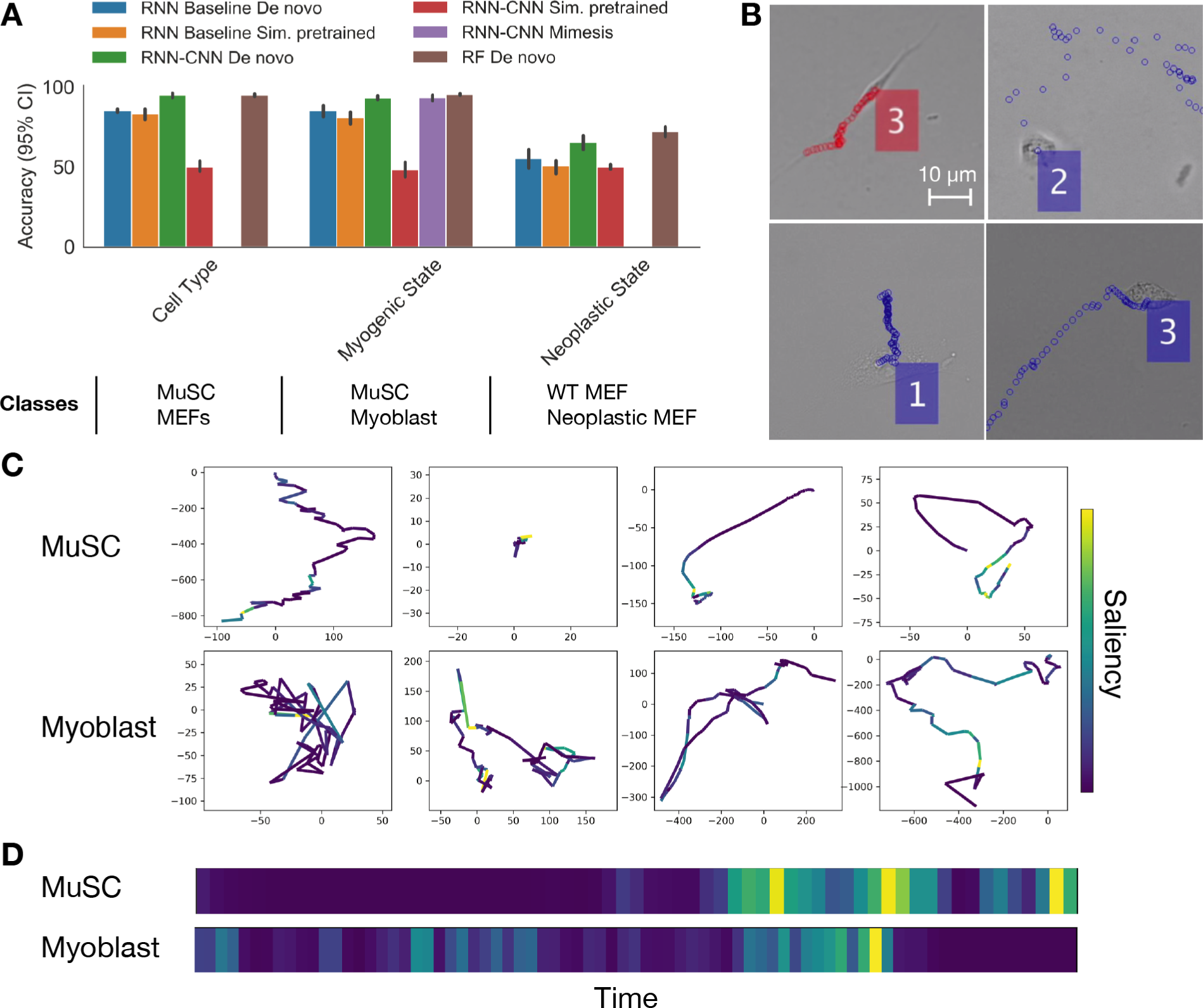
RNNs can discriminate between different cell types and cell states based on motility. (A) Prediction accuracy (5-fold CV, mean ±95%) CI) for RNN models and a heuristic baseline trained to classify experimentally measured cell motility. Models pretrained with weights from a simulated motion classifier or a cell mimetic motion classifier are indicated in the legend. RNN-CNN models with mimetic pretraining were only trained for myogenic state classification and therefore data are not presented for other tasks. (B) Representative images of different cell types, from top left to bottom right: MuSC, myoblast, neoplastic MEF, and wild-type MEF. Colored markers indicate the cell’s path along the substrate over time. (C) Saliency analysis of an RNN classifier for myogenic activation states overlaid with cell tracks in the Cartesian plane. Large displacements in the myoblast tracks are highly salient, while in MuSC tracks sharp turns and characteristic “U-turns” are salient. (D) Saliency analysis presented as a heatmap for representative myoblast and MuSC motility tracks.

Transfer learning using simulation classification as a pretraining task decreased cell type classification performance for both baseline RNNs and RNNs with convolutional layers (Fig. 4A). This result suggests that the task of classifying simulated motility is too distinct from the challenge presented by experimentally measured cell motility to provide benefit. This suggests that more realistic simulations may yield the desired performance improvements.

### D. RNNs provide discriminative power between stem cell activation states

To determine if RNNs can distinguish between more nuanced differences in cell state, RNN classifiers were trained to discriminate between myogenic activation states (MuSCs and myoblasts). Training was performed on *n* = 334 MuSCs and myoblasts per class using 5-fold cross-validation. As with the cell type classification experiments, 10% of each training fold was used for testing after each epoch and model selection by early stopping.

Mean validation accuracy reached ∼ 85% for RNN baseline models, ∼ 93% for RNN models with convolutional layers, and ∼ 95% for our heuristic baseline (Fig. 4A). As with cell type classification, the increase in performance provided by the addition of convolutional layers to the RNN baseline model is significant (*p* < 0.05, *t*-test). Again similar to cell type classification experiments, pre-training using simulated motility tracks decreased rather than increased RNN performance. These results demonstrate that RNN models can discriminate between stem cell activation states based on motility alone, even with small data sets. The RNN models perform comparably to hand-engineered features paired with a Random Forest classifier on this task.

RNN classifiers were also trained in the same manner to discriminate between wild-type and neoplastic MEFs with transfer learning from the simulated motion classifier. Training was performed on *n* = 250 samples per class using 5-fold cross-validation. Testing data was held out from each training fold as before. RNN models failed to achieve validation accuracy greather than ∼ 65% on this task (Fig. 4A). The baseline heuristic model performed at 72% accuracy, significantly outperforming the RNN models.

The more nuanced phenotypic difference between wild-type and neoplastic MEFs may be an inherently more challenging classification problem. The small available sample size likely compounds this difficulty and exacerbates the RNN classifiers’ poor performance. This experiment highlights the fact that hand-engineered features may provide superior performance on some tasks.

To develop an understanding of how the RNN models were discriminating MuSC and myoblast motility, we performed saliency analysis to visualize the relevant aspects of our input tracks for classification [50]. The basic principle of saliency analysis is to identify the regions of a given input that have the most influence on whether or not the input is considered part of a particular class. This influence of input regions on a particular classification decision is estimated by the magnitude of the gradient on the input with respect to the class score (see Methods). Here, we find the temporal regions of cell motility tracks that provide the highest signal for the network to classify that input as either a MuSC or a myoblast (Fig. 4C, D). Qualitatively, it appears that large displacements in myoblast tracks provide strong gradients for the myoblast class, while tight turns and reversals of direction in MuSC tracks provide strong gradients for that class.

Saliency analysis of this form offers a unique window for interpreting motility classification models. While interpretability techniques are available for hand-engineered features, it is often difficult to dissect nuanced interactions between features that may be causal for classification decisions for a given sample. Saliency analysis offers the ability to highlight input regions, rather than pre-computed features, which lead to a classification result. These highlighted regions of input may be easier to interpret than a set of most influential features in some instances.

### E. Cell mimetic pretraining

Transfer learning from RNN classifiers trained on random walk, Levy flight, and fractional Brownian motion simulations failed to improve classification performance on experimentally measured cell motility tasks. We reasoned that this may be due to the drastic differences between the behavior of these simulations and real cells. To remedy this, we attempted to generate simulated data that more accurately reflected real cell motility to enhance pre-training efficacy. For a set of real cell motility data, we measure the displacements and turning behavior of each cell. Displacements are measured simply as the Euclidean distance between each set of sequential timepoints. The turning direction at a point *t_i_* is determined as the angle between the vectors that connect points *t_i−_*_1_ to *t_i_* and *t_i_* to *t_i_*_+1_.

Cells are decomposed into a set of *k* clusters by *k*-means clustering on a set of parameters measured from these displacement and turn angle distributions. The number of clusters *k* = 5 was chosen empirically to capture the diversity of the cell phenotypes while still leaving non-trivial numbers of cells in each cluster. For each cluster, a bounded Johnson distribution is fit to the aggregate distribution of displacements and the aggregate distribution of turn angles. Simulated samples are generated by randomly sampling displacement magnitudes and turn angles from the fitted Johnson distributions for *T* time steps. To represent a population of cells, the proportion of simulations generated from each cluster is equivalent to the cluster’s prevalence in the original cell data. This approach may be conceptually likened to the bag-of-words model [63], in which *k*-means clustering is used to decompose features into a representative “vocabulary.” By sampling from each of *k* clusters proportionally, we aim to capture and simulate heterogeneous phenotypes within a cell population, rather than simply reproducing a single averaged phenotype that may not be representative of any true cell phenotype.

We generated “cell mimetic” simulations for MuSCs and myoblasts by the above method, with *n* = 15, 000 simulated samples for each of the two activation states. RNN classifiers were pretrained by classifying between the two simulated data sets, reaching ∼ 99% validation accuracy. The weights from this pretrained network were used to initialize an RNN classifier for the myogenic activation task outlined above. Mean validation accuracies were effectively unchanged at ∼ 93% for the RNN model with convolutional layers. These results indicate that pre-training on “mimetic” simulations has little effect on final classification accuracy, and that transfer learning from simulations to experimentally measured data is not advantageous for the tasks explored here.

### F. Autoencoders allow unsupervised learning of representations in motion feature space

Results up to this point indicate that supervised classification of different cell motility phenotypes using RNN models is effective and comparable in performance to classification with hand-engineered features. However, in the analysis of motility data, supervised classification data is not always available. For instance, to explore the heterogeneity of types in a given population, there is no obvious method to generate supervised classification data that may be used to learn relevant feature kernels by optimization of a standard classification loss function. This would also be an issue in the identification of heterogeneous motility behaviors in patient biopsy samples, in which the distinguishing features are not known *a priori*. Training neural networks as autoencoders in an unsupervised fashion has been used in other contexts to learn relevant feature kernels where no obvious classification problem is present [28], [29].

In addition to learning useful features, autoencoders can be used to map high-dimensional inputs to a low dimensional latent space for the identification of subpopulations. This use of autoencoders has proved successful when applied to RNA-sequencing data [64], [65] for cell type and cell state identification. Similarly, the latent dimensions of an autoencoder may serve to segregate cell states from cell motility data in an unsupervised fashion. To determine if autoencoders can learn unsupervised representations of cell motility, we trained RNN autoencoders on our multi-channel time series representations of cell motility.

An RNN autoencoder was formulated by using the same initial convolutional layers and LSTM unit as in our classification architecture. Following the LSTM unit, we again use a set of three fully-connected layers paired with ReLU activations.

We set the central fully-connected layer to use *n* = 16 hidden units, corresponding to 16 latent dimensions. The number of latent dimensions was chosen empirically and does not represent the result of a rigorous hyperparameter search. Following the fully connected layers, we utilize upsampling and convolutional layers to decode the latent representation back to the input dimensions (Fig. 1B).

RNN autoencoders were trained similarly to classification models, but the regularization strength was increased to *λ* = 10^−5^. Regularization strength was increased after over-fitting was observed empirically. The *Adadelta* optimizer was used with learning rate *ϵ* = 0.1 [66]. The alternative optimizer was chosen empirically based on the time to convergence we observed in early training experiments.

To determine if the autoencoder could learn features from simulations in an unsupervised manner, the model was trained on *n* = 15, 000 samples of each class for three types of simulated motion (random walk, Levy flight, fractional Brownian motion) using 5-fold cross validation. Mean squared error was used to as a loss function. Models consistently converged for each training fold (MSE = 410 ± 11, pixel units). RNN autoencoder outputs consistently failed to capture the full extent of a the input motility track, seeming instead to capture a vague notion of the motility location and extent (Fig. 5A).

**Fig. 5.**
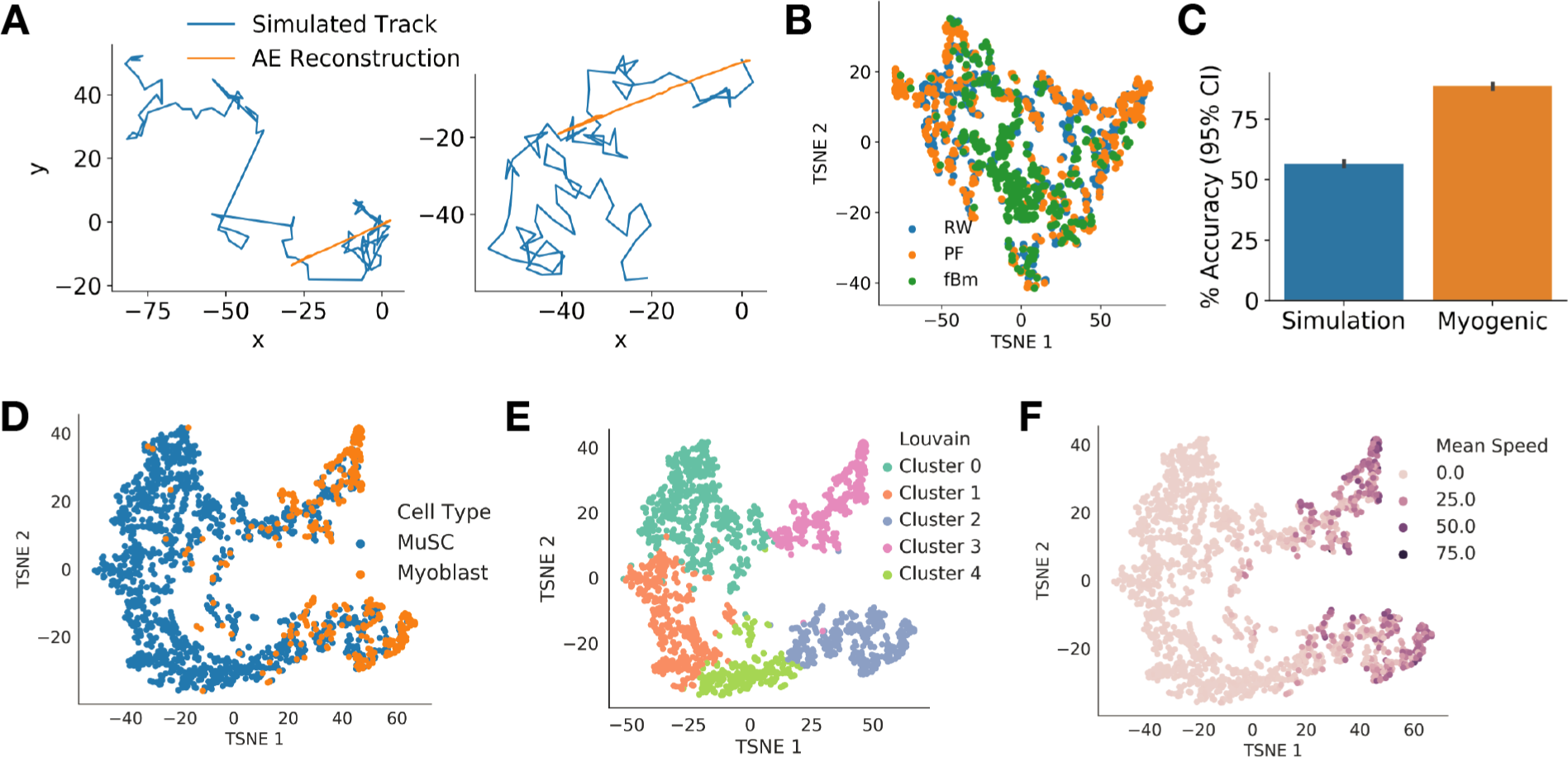
RNN autoencoders can learn representations of cell motility in an unsupervised manner. (A) Sample input and (B) output to an RNN autoencoder. (C) Accuracy of Random Forest classifiers trained to discriminate simulations or myogenic states based on RNN autoencoder latent features. (D) t-SNE visualization of the latent space learned by an RNN autoencoder trained on MuSC and myoblast cell motility tracks. The model segregates the different states of myogenic activation and finds a continuous progression from the MuSC to myoblast state. (E) Louvain community detection identifies multiple myogenic sub-states. Myoblasts are largely segregated into two clusters, shared with more myoblast-like MuSCs. (F) Mean speed features overlaid on the RNN latent space. The autoencoder is able to capture properties of motion we might extract heuristically.

Do the latent dimensions capture information about the inputs, even if the reconstructions are poor? To answer this question, we utilized the output of the autoencoders’ central layer (the encoded representation) as features to classify the input classes for each simulation. A Random Forest classifier was trained to distinguish the simulation classes from these features. Random Forests trained on autoencoder features achieved ∼ 56.4% accuracy on this 3 class problem. This indicates that the features learned by these autoencoders contain some information about the three simulated classes, but are less useful for classification that the heuristic features we define as a baseline (Fig. 5C).

The encoded representation of an autoencoder may also be interpreted as a latent space where clusters within the data may be identified. Visualizing the latent space of our autoencoder trained on simulated tracks using t-SNE [67] reveals that despite the poor quality of reconstructions, the latent dimensions largely segregate fractional Brownian motion (fBm) simulations from random walks (RW) and Levy flights (PF), though the latter two are intermixed (Fig. 5B).

It is possible that a similar examination of the latent space in an autoencoder trained on experimentally observed cell tracks will reveal structure within the data. To determine if our RNN autoencoder models were capable of learning latent spaces that segregate cell types and cell states, we trained an autoencoder on MuSC *n* = 1407 and myoblast *n* = 334 tracks (length *T* = 60) using 5-fold cross-validation. The models consistently converged on each training fold (MSE = 8574 ± 830, pixel units). Training a Random Forest classifier to discriminate MuSCs and myoblasts based on the 16 latent dimensions of this model, as for the simulated motility autoencoder above, yields a prediction accuracy ∼ 88%. Again, this performance is inferior to that achieved using hand-engineered features, but does demonstrate that the latent dimensions capture variation between myogenic cell states (Fig. 3C).

Examining a t-SNE visualization of the latent space, it is evident that the latent dimensions capture variation between MuSCs and myoblasts, segregating myoblasts into one region of the latent space (Fig. 5D). Interestingly, the latent space does not separate the two cell types discretely, but rather presents a continuum between the MuSC and myoblast phenotypes, such that some MuSCs are similar to myoblasts, while others are distinct. This reflects the underlying biology, in which MuSCs transition over time from a quiescent state into the activated myoblast state [68], [69].

We apply Louvain community detection [70], as commonly employed in the field of single cell RNA-sequencing [71], to identify subpopulations within the latent space. Community detection identifies five subpopulations, capturing the majority of myoblasts in two of the five clusters (Fig. 5E). It is interesting that myoblasts are segregated into two non-overlapping clusters, suggesting two discrete states of myoblast motility. MuSC captured within the myoblast dominated clusters (2 and 3) may be more activated than counterparts in cluster 1, while cluster 4 may represent an intermediary state. Cluster identification within these autoencoder latent spaces is a promising approach to reveal substructure within cell populations. To determine if our autoencoder model was learning some heuristic features, we visualized the mean speed of each track in the latent space (Fig. 5F). It is readily apparent that the model learns to discriminate cells with different mean speeds, indicating that the model is capable of capturing some of the features we may have hand-engineered in a completely unsupervised manner.

Collectively, these results demonstrate that RNN autoen-coders are capable of distinguishing cell types and revealing heterogeneous substructure from cell motility observations in an entirely unsupervised manner. The continuum of cell states between MuSCs and myoblasts suggested by the autoencoder latent space is consistent with known biology and suggestive of trajectories cells traverse through behavioral space. Application of these methods to other cell types may similarly suggest trajectories of cell state transition.

### G. RNNs predict muscle stem cell motility

Tracking individual cells in timelapse microscopy experiments is a difficult multi-object tracking problem [72]. Popular tracking methods utilize a motion model to predict cell motility in advance of the next frame to improve tracking performance [73]. This motion prediction is especially useful in the event of “missed detections,” where a cell is not detected or segmented for a given set of frames but is detected later on. The most common motion models employed are based on linear kinematics, with Kalman filters serving as a popular choice [52]. Linear kinematics assume that particles possess an inertia and tendency to continue moving in the same direction as previously observed. These assumptions are equivalent to assuming objects exhibit ballistic motion.

However, cell motion does not adhere to ballistic assumptions in all cell types, with myogenic cells being an excellent example of such a system. In myogenic cells, a motile cell often makes a few rapid displacements punctuated by stopping periods or direction changes which violate intuitions about the inertia of moving objects. A motion model specifically tailored to the cell type of interest may therefore be useful to improve tracking performance, but such specific tailoring would require a prior knowledge of the very motion features that the live cell experiment is designed to analyze. Some way to tailor prediction models on the fly could help solve this problem.

Recurrent neural networks have been effectively utilized for sequence prediction in multiple fields [74], [75], [76], [77]. We adapted the convolutional RNN autoencoder model to a sequence prediction model by removing the pooling layers and fully-connected layers and altering the number of nodes in the central LSTM layer (Fig. 6C, see Methods). As a prediction task, we trained the RNN prediction model on *τ_in_* = 20 time steps of motion and predicted *τ_out_* = 10 time steps into the future. As a data set, we split MuSC motion paths into subpaths of length *τ_total_* = 30 for a total of *n* = 8, 676 paths.

**Fig. 6.**
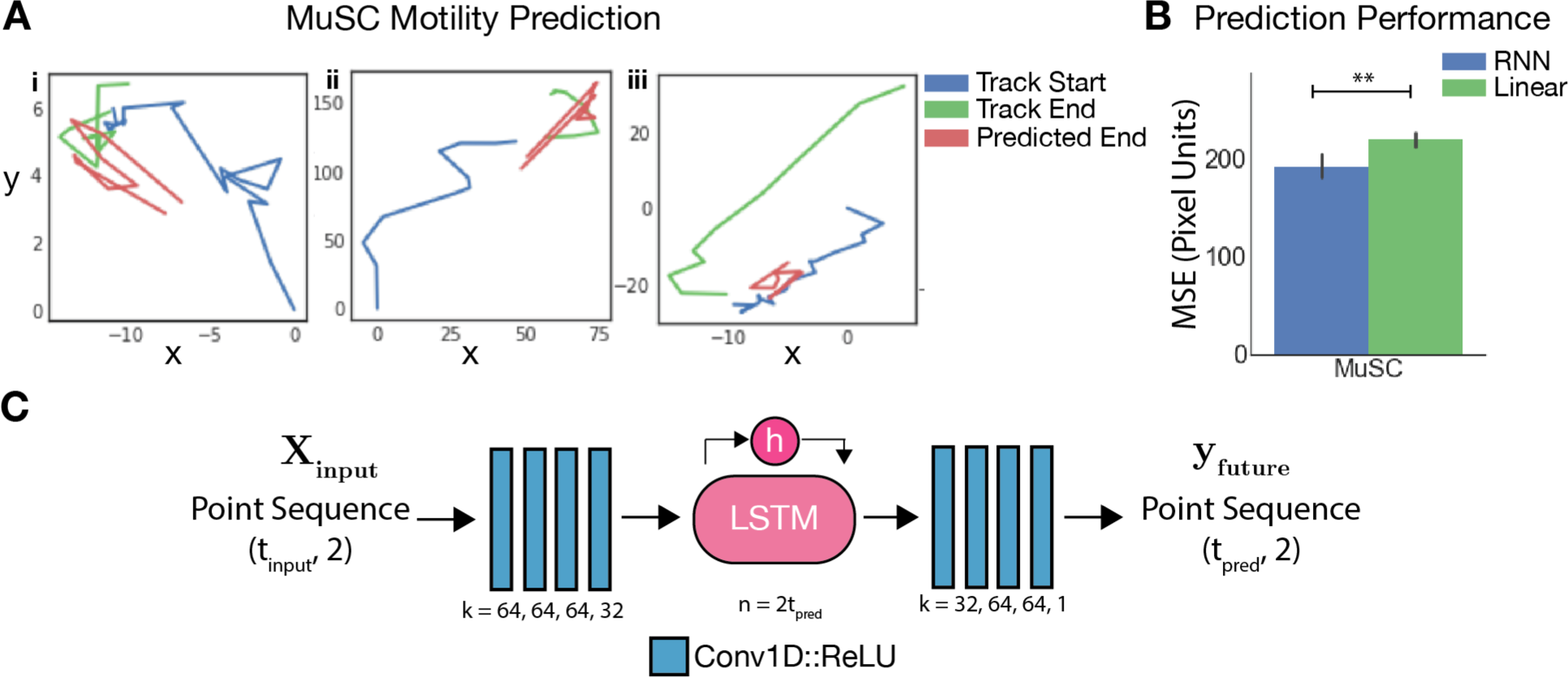
RNN models predict MuSC motility more effectively than linear kinematic models. (A) Representative samples of MuSC tracks used for prediction with predicted track endings and true track endings. (B) Performance of the RNN motion prediction model relative to a linear kinematic baseline model, determined as the mean squared error between ground truth and predicted track endings. (\*\**t*-test *p* < 0.001) (C) RNN motility prediction architecture, where *k* is the number of kernels in each 1D convolutional layer and *n* is the number of units in the LSTM. All convolutional layers except the final layer are paired with a ReLU activation.

A validation set of 10% of all tracks was held out, and the remaining tracks were split with 80% used for training and 20% used for testing. Mean squared error between the predicted path and the ground truth path was used as the loss function and the *Adam* optimizer was used with learning rate *ϵ* = 0.001. No regularization was used when training motility prediction models.

As a baseline for comparison, we performed a simple kinematic prediction of MuSC paths that assumes persistence of the velocity from preceding time points. The velocity for prediction was obtained by averaging instantaneous velocity for *τ* = 15 time points prior to the track terminus, where *τ* was optimized by parameter search. This baseline model leads to an average mean squared error (MSE) in pixel units of ∼ 220 (30 train/test splits). The RNN prediction model by comparison produces a significantly lower MSE of ∼ 192 (pixel units) (*t*-test *p* < 0.001), indicating that the RNN model is a superior motion predictor in the MuSC context (Fig. 6B). Representative track endings (length *τ_out_* = 10) produced by the RNN prediction model are displayed alongside the preceding track beginnings (length *τ_in_* = 20) and the ground truth track endings (Fig. 6A).

In most cases the motion prediction reasonably approximates the cell’s ground truth direction, but does not closely mirror the exact path (Fig. 6A, inset **i** and **ii**). In some events, the RNN model fails to predict even the correct direction of motion (Fig. 6A, inset **iii**). We performed the same experiment with mimetic myoblast simulations using *n* = 10^5^ total samples, holding *n* = 5000 samples for validation. Similar to the MuSC results, RNN motion predictors achieved a markedly lower tMSE of 1195, relative to the baseline kinematic model MSE of ∼ 9797 (*t*-test *p* < 0.001).

These results indicate that convolutional RNN models can be effective cell motility prediction models and are superior to simple linear kinematic approaches in some real world circumstances. RNN motility prediction models may therefore offer a scalable way to fit a uniquely tailored motion model to specific cell biology contexts. These cell-context specific RNN motility predictors may be useful to improve multi-cell tracking performance, which remains a difficult problem in the field. Addressing challenges with cell tracking is a problem well suited for the type of learned modeling approach we present, as different cell types can exhibit highly different motility behaviors, making the *a priori* design of a universal motion prediction model difficult.

## IV. Conclusion

Deep neural networks enable representation learning, or learning of features relevant for the description of a feature space. By representing cell motility as a multichannel time series, we show that RNNs with convolutional feature extractors may be applied as an effective analytical tool. Our results demonstrate that these models are capable of discriminating between simulated models of motion and multiple types of experimentally measured cell motility behaviors, though these models are still inferior to hand-engineered features paired with random forest classifiers. In our experimentally measured cell motility data, we find that RNN models effectively discriminate between different cell types, and different states of myogenic progenitor activation. We also find that RNN autoencoders can learn latent space that distinguish between cell states and suggest cell state transition trajectories in an unsupervised fashion.

Adapting the convolutional RNN autoencoder for motility prediction, we find that the RNN model is more effective at predicting MuSC motility than a standard linear model. Such prediction models may be useful for cell tracking. While we apply the methods described here to cell biology, there is no conceptual limitation that prevents application to other fields where discrimination based on motion recordings is desired. In the field of cell biology, analysis of motility with deep neural networks may allow for useful insights to be gathered in contexts where relevant features are non-obvious or laborious to construct.

## Acknowledgment

This work was funded by NSF grant MCB-1515456 to W.F.M., NIH grants AR060868 and AR061002 to A.S.B., and a PhRMA Foundation fellowship to J.C.K. This material is based upon work supported by the National Science Foundation Graduate Research Fellowship under Grant No. 1650113 to J.C.K. J.C.K and W.F.M. were also supported by the NSF Center for Cellular Construction, NSF Grant No. DBI-1548297. We gratefully acknowledge the support of NVIDIA Corporation with the donation of a Titan Xp GPU used for this research. We would like to thank Morgan Truitt and Davide Ruggero for providing neoplastic MEF cells.

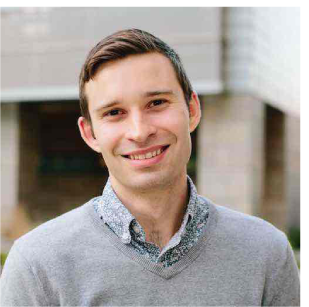

**Jacob C. Kimmel** Jacob Kimmel is a Data Scientist at Calico Life Sciences, where he uses quantitative biology methods to study the biology of aging. Jacob previously earned a PhD in Developmental & Stem Cell Biology at the University of California, San Francisco. His research focuses on the application of machine learning methods to infer cellular phenotypes from cell behaviors and quantification of cell state dynamics.

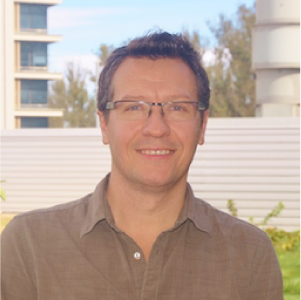

**Andrew S. Brack** Andrew Brack has been trained as a molecular and developmental biologist. In his PhD work, working under Prof. Malcolm Irving (FRS), Andrew used a combination of biophysical and molecular biology approaches to understand the structural mechanisms that control skeletal muscle contraction. Following his PhD, Andrew performed postdoctoral research at Kings College London with Dr. Simon Hughes and then at Stanford University with Dr. Tom Rando, where he studied skeletal muscle stem cell regulation. He is currently an Associate Professor of Orthopedic Surgery at UCSF, where his research focuses on understanding the cellular and molecular mechanism that control skeletal muscle regeneration.

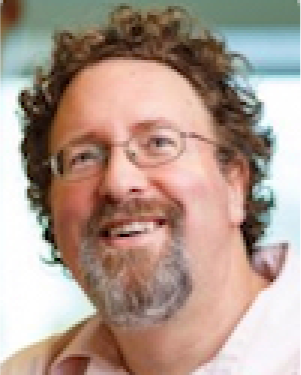

**Wallace F. Marshall** After undergraduate training in Electrical Engineering at SUNY Stony Brook, Wallace Marshall obtained his Ph.D. in Biochemistry at UCSF, followed by postdoctoral research in cell biology at Yale University. He is currently professor of Biochemistry and Biophysics at UCSF, where his research focuses on how cells solve engineering problems related to establishment of cellular geometry, including analysis of organelle size control systems, mechanisms for regeneration in single cells, and cellular decision making.

